# Effect of pulsed light on postharvest disease control-related metabolomic variation in melon (*Cucumis melo*) artificially inoculated with *Fusarium pallidoroseum*

**DOI:** 10.1101/698407

**Authors:** Francisco Oiram Filho, Ebenézer de Oliveira Silva, Mônica Maria de Almeida Lopes, Paulo Riceli Vasconselos Ribeiro, Andréia Hansen Oster, Jhonyson Arruda Carvalho Guedes, Patrícia do Nascimento Bordallo, Guilherme Julião Zocolo

**Author notes:** Corresponding author (GJZ).

## Abstract

Pulsed light, as a postharvest technology, is an alternative to traditional fungicides, and can be used on a wide variety of fruit and vegetables for sanitization or pathogen control. In addition to these applications, other effects also are detected in vegetal cells, including changes in metabolism and production of secondary metabolites, which directly affect disease control response mechanisms. This study aimed to evaluate the possible applications of pulsed ultraviolet light in controlling postharvest rot, mainly caused by *Fusarium pallidoroseum* in yellow melon ‘Goldex’, *in natura*, and its implications in the disease control as a function of metabolomic expression to effect fungicidal or fungistatic. The dose of pulsed light (PL) that inhibited *F. pallidoroseum* growth in melons (*Cucumis* melo var. Spanish) was 9 KJ m^-2^. Ultra performance liquid chromatography (UPLC) coupled to a quadrupole time-of-flight (QTOF) mass analyzer identified 12 compounds based on the MS/MS fragmentation patterns. Chemometric analysis by Principal Components Analysis (PCA) and Orthogonal Partial Least Squared Discriminant Analysis (OPLS-DA and S-plot) were used to evaluate the changes in fruit metabolism. PL technology provided protection against postharvest disease in melons, directly inhibiting the growth of *F. pallidoroseum* through upregulation of specific fruit biomarkers such as pipecolic acid (**11**), saponarin (**7**), and orientin (**3**), which acted as major markers for the defense system against pathogens. PL can thus be proposed as a postharvest technology to avoid chemical fungicides and may be applied to reduce the decay of melon quality during its export and storage.

## Introduction

The melon (*Cucumis melo* L.) is a tropical fruit and is an economically crucial part of Brazilian exports. However, the major fragilities in the postharvest chain of melon are incidences of postharvest pathologies, particularly the rot caused by *Fusarium pallidoroseum*, which is responsible for postharvest losses of melons. The melon is a ground plant whose fruit is in contact with soil, thus facilitating its contamination with *F. pallidoroseum* [1]. *F. pallidoroseum* is a widespread and common species in tropical, subtropical, and Mediterranean areas, often isolated from plants with complex diseases; it is also known to be toxigenic [2].

Fungal disease from genus *Fusarium* in fruit is traditionally controlled by application of synthetic fungicides that leave chemical residues, which could be harmful to the consumer, and also encourage the development of fungicide-resistant strains of fungal pathogens. To overcome these challenges, several alternatives or integrative approaches, including physical methods, are imperative to develop fruit with increased natural defense responses through the resistance mechanisms induced by abiotic stress [3].

Among these strategies, application of technologies using light as an abiotic factor aiming to promote a regulatory and signaling role in the developmental and metabolic processes of plants is encouraged [4]. Ultraviolet light (UV-C) is a continuous radiation that is widely reported in the induction of resistance mechanisms and control of postharvest diseases, thus extending the shelf-life of fruit and vegetables [5, 6]. However, studies involving pulsed light (PL) as radiation from the perspective of inducing resistance mechanisms are still discreet [7].

PL is a new, non-thermal technology, where a lamp containing an inert gas such as xenon emits high-frequency radiation pulses between wavelengths of 180 to 1100 nm [8]. This technology is used to sanitize fruit and vegetable surfaces by acting on microorganism cells and breaking and altering their DNA sequences, thus inhibiting the pathogen reproductive capacity [9]. Moreover, the use of PL as a physical method can stimulate the production of phytochemicals in plant tissues to minimize the possible deleterious effects caused by radiation [10].

Metabolomics has been emphasized within the “omics” sciences, and efficiently evaluates a large part of the metabolites of an organism both quantitatively and qualitatively, at a given time and in a specific situation [11]. Metabolomics can identify a change in the concentration of compounds involved in primary metabolism or production of secondary metabolites. These metabolic alterations may arise from cellular lesions, metabolic adjustments to restore cellular homeostasis, or synthesis and/or accumulation of metabolites resulting from cellular pathways [12].

The major objective of this study was to investigate the fungicidal or fungistatic effect to *Fusarium pallidoroseum* in the melon cv. Spanish by PL treatment using ultra-performance liquid chromatography-quadrupole-time-of-flight mass spectrometry (UPLC-QTOF-MS^E^) and chemometric tools to identify possible biomarkers associated with metabolic changes in the defense mechanism.

## Material and methods

### Chemical compounds

Acetonitrile (PubChem CID: 6432), formic acid (PubChem CID: 284), methanol (PubChem CID: 887), sodium hypochlorite (PubChem CID: 23665760), MilliQ water (PubChem CID: 962), liquid nitrogen (PubChem CID: 947).

### Plant material

Melons (*C. melo* var. Spanish) were obtained at maturity stage (10–12 °Brix) from a commercial growing field of Norfruit Northeast Fruit, located at Mossoro-RN, Brazil (04”54’9,4”S and 37”21’59,9”W). Mature melons were surface disinfected with 200 μg L^-1^ sodium hypochlorite solution for 2 min, rinsed, and allowed to drain. The melons were then artificially inoculated.

### Fruit inoculation

*F. pallidoroseum* spore suspension was produced from pure colonies of the fungus at a concentration of 10^6^ spores mL^-1^. Melons were inoculated near the peduncles and adjacent regions with 100 µL inoculums of *F. pallidoroseum* (n = 5 inoculants). After inoculation, the melons were subjected to PL treatment.

### Disease control by PL treatment

After 12 h there is already pathogen effect [13] on fruits inoculated with *F. pallidoroseum*. Then, the fruits were irradiated in a PL chamber (XeMaticA-2LXL model, SteriBeam® GmbH, Germany) equipped with two xenon flash lamps and Teflon® transparent supports that allowed the melons to be uniformly exposed over 360° by both lamps with uniformity at distance of 0.07 m between fruit and lamps. The lamps produced short-time pulses of 0.3 µs, delivering broad-spectrum white light (200-1100 nm) with approximately 15–20 % of UV-C, according to the system built-in photodiode readings. Evaluation of disease control against *F. pallidoroseum* was carried out by dose screening, as follows: 0 (non-treated with PL), 6, 9, and 12 KJ m^-2^, according to the standard limits proposed [14]. After PL-screening, the inoculated and PL-treated melons were incubated in a box protected from light for 48 h at 28 °C ± 1 °C with relative humidity of 92 % to ensure optimal conditions of mycelium growth, and then stored at room temperature for 21 d. After this time, the occurrence of a severe degree of fungal disease was analyzed by measuring the radial lesion diameter in the inoculum region using a digital pachymeter (Digimess®, São Paulo, Brazil), and expressing it in meters (m) and percentage of disease incidence (%). These analyses were performed using randomized design, each treatment comprised 8 replicates and data were subjected to analysis of variance (ANOVA) followed by Tukey’s test at 5% probability.

### Disease control promoted by suitable PL-dose against *F. pallidoroseum*

After PL-screening, an additional experiment was conducted using the suitable PL-dose capable of controlling fungal disease in the fruit. To evaluate disease control, melons were inoculated with *F. pallidoroseum* under the same inoculation conditions described in Fruit inoculation section. Inoculated melons were then irradiated with a PL-dose of 9 KJ m^-2^ in the same instrumental conditions described in Disease control by PL treatment section. The control group included inoculated and non-inoculated melons. After 96 h, biological triplicates of each treatment were subjected to extraction processes for injection into a UPLC system.

### Extract preparation

Extracts were obtained Moore, Farrant (15] with modifications. The pellets were extracted from the different adjacent regions (0.005 m) to the inoculum with thickness approximately of rind melon. Peel melon powder (1.00 g) was resuspended in 4 mL of MeOH/H_2_O (7:3 *v/v*). The homogenate was then subjected to sonication (Ultracelaner 1450, Unique®, Brazil) for 30 min, followed by centrifugation at 6,000 × *g*, for 5 min. The pellet was re-extracted twice using 3 mL of MeOH/H_2_O (7:3 *v/v*) using the same conditions of ultrasound and centrifugation. The supernatant was filtered through a 0.22 μm polytetrafluoroethylene membrane (PTFE) (Biotechla®, Bulgaria) and injected directly into a UPLC system.

### Chromatographic analysis by UPLC-QTOF-MS^E^

The analyses were accomplished on an Acquity UPLC (Waters, USA) system coupled to a Xevo Quadrupole and Time-of-Flight mass spectrometer (Q-TOF, Waters). Separations were performed on a C18 column (Waters Acquity® UPLC C18 - 150 mm × 2.1 mm, 1.7 μm). For metabolic fingerprinting, a 2-μL aliquot of phenolic extract was subjected to UPLC using an exploratory gradient with the mobile phase composed of deionized water (A) and acetonitrile (B), both containing formic acid (0.1% *v/v*). The phenolic extracts from melons were subjected to exploratory gradient as follows: 2–95% for 15 min, with a flow rate of 500 µL min^-1^. Ionization was performed with an electrospray ionization source in negative mode ESI, acquired in the range of 110–1200 Da. The optimized instrumental parameters were as follows: capillary voltage at −2800 V, cone voltage at −40 V, source temperature at 120°C, desolvation temperature at 330°C, flow cone gas at 20 L h^-1^, desolvation gas flow at 600 L h^-1^, and the microchannel voltage of the plate (MCP) - detector at −1900 V. The mode of acquisition was MS^E^ and system was controlled using MassLynx 4.1 software (Waters Corporation). The extracts were injected in triplicate.

### Statistical analysis

The UPLC-MS data were processed using MassLynx® software (Waters Co., Milford, MA, USA), under the following conditions: retention time deviation tolerance ± 0.05 min, mass range 110–1200 Da (accurate mass tolerance of ± 0.05 Da) and noise elimination level 5. For structural identification of metabolites, molecular formulas were considered and the values for *m/z* were obtained from high-resolution spectra observed in the chromatogram at higher intensity. The relative error is given in ppm for each formula. Margins of error less than 10 ppm are considered for studies in MS/MS.

The structural proposals of molecules were performed using MS/MS data through the establishment of rational fragmentation patterns reported in literature [16-19]. A list of identities of peaks was created using the retention time (rt) and error (*m/z*). For unidentified peaks, all possible molecular formulas were derived (elements C, H, N, and O, with the tolerance of 10 ppm, at least 2 carbon atoms) using the elemental composition tools available in MassLynx® software.

The UPLC-MS data analyzed by chemometrics were processed using Markerlynx® software for Principal Components Analysis (PCA) and Orthogonal Partial Least Squared Discriminant Analysis (OPLS-DA and S-plot). S-plots were obtained via OPLS-DA analysis to determine potential biomarkers that significantly contributed to the difference among groups [20-23].

## Results and discussion

### The growth of *F. pallidoroseum* under PL treatment

The lesion diameter of fungal infection (Fig 1) and percentage of disease incidence (data not shown) were analyzed to determine the suitable PL-dose of treatment for the control or inhibition of *F. pallidoroseum* in melon fruit. The growth of the pathogen on melon fruit inoculated with *P. pallidoroseum* was strongly inhibited by PL treatment (Fig 1). This is corroborated as shown in Fig 1, by a mean lesion diameter of 0.013 m, with a 100% incidence of fungal disease in melons without PL radiation.

**Fig 1.**
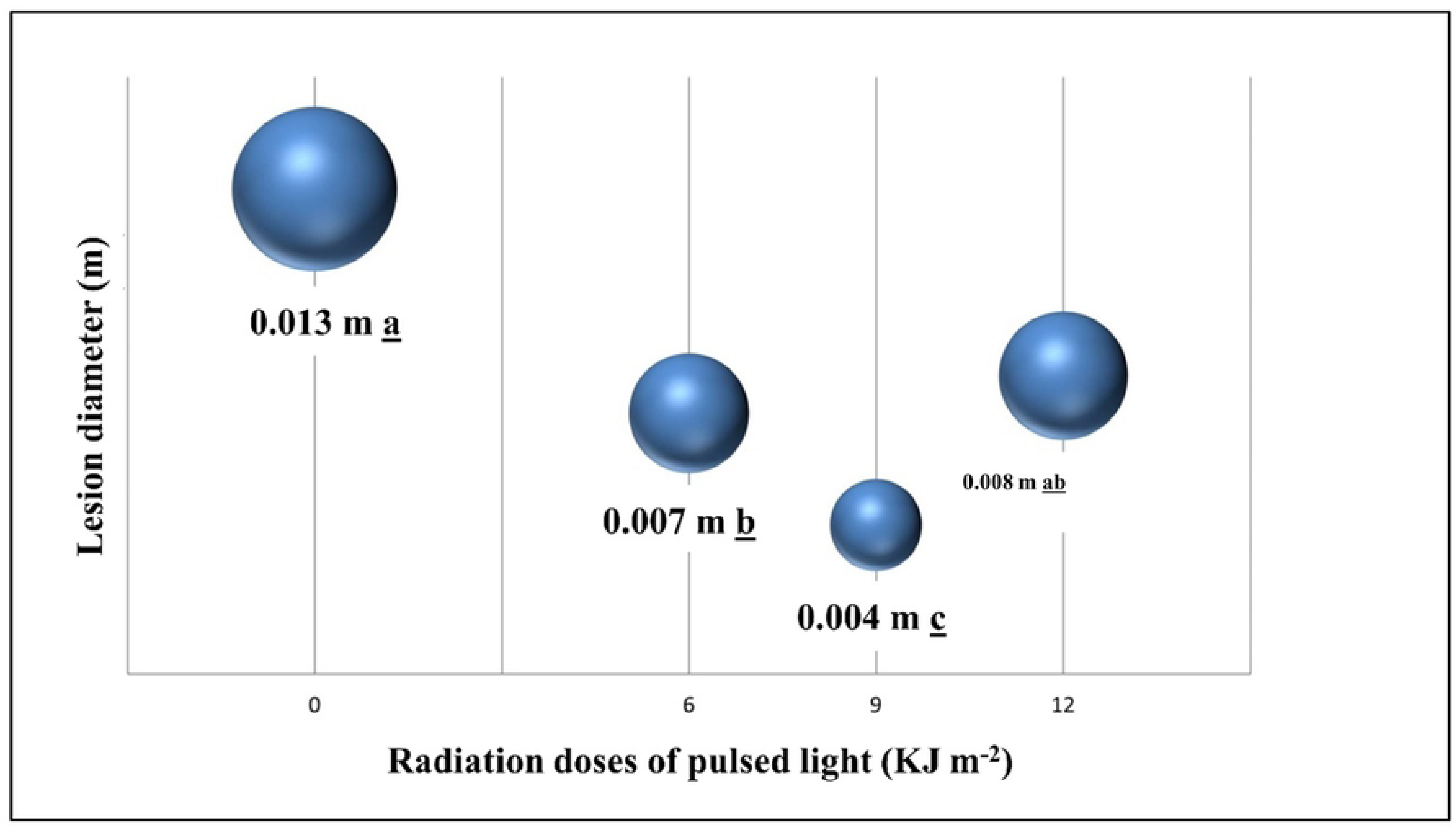
Screening of PL doses to evaluate the lesion diameter (m) in melon inoculated with *F. pallidoroseum*. Mean values followed by the same small letter did not differ significantly between PL-treatments, by Tukey’s test at 5% probability.

Disease progression in the PL-treated group that received 6 KJ m^-2^ was significantly lower than that in the untreated group, with a significant reduction in lesion diameter (0.007 m) (Fig 1) and 62.5% incidence of fungal disease. Moreover, a dose of 9 KJ m^-2^ was more effective than all the doses employed and was associated with a low mean lesion diameter of 0.004 m, (Fig 1), and 33.3% disease incidence by *F. pallidoroseum*. The hormesis concept was used to explain the behavior with a dose radiation of 9 KJ m^-2^.

Hormesis is a phenomenon in which low levels of potentially damaging radiation elicit beneficial responses, i.e. physiological stimulation of beneficial responses in plants by low levels of stressors that otherwise elicit harmful responses. Hormetic doses of ultraviolet light (UV-C) radiation are involved in plant susceptibility towards diseases, and are capable of elicit plant resistance mechanisms including production of anti-fungal compounds such as specific phenolics including phytoalexins that act both as light quenchers that absorb damaging wavelengths of light, and antioxidants that prevent reactive oxygen species (ROS) mediated cellular damage [7, 24, 25]. The UV-C radiation might also have fungistatic effect promoted by phenolic compounds which plays a barrier role physically against pathogens attacks and the same barrier reduce water and nutrients diffusion important to the growth pathogen [26].

However, a recent study compared the application of low-intensity UV-C and high-intensity PL sources as elicitors of hormesis in tomato fruit (*Solanum lycopersicum* cv. Mecano) [7]. Curiously, these authors showed that postharvest hormetic treatment with 16 pulses of PL (7.4 KJ m^-2^) with a spectral range (240 – 1050 nm) significantly delayed ripening along with inducing disease resistance to *Botrytis cinerea* in tomato fruit with 41.7% reduction in disease progression compared to 38.1% reduction with the conventional low-intensity UV-C (254 nm) at 0.37 KJ m^-2^. Thus, according to the authors, PL treatment, although rich in UV-C (broader spectral output), elicited the same pathways or responses as hormesis induced by conventional low UV sources (narrower spectral range), making the PL-treatment more commercially attractive, as this treatment allows a substantial reduction in treatment time from seconds to microseconds.

The last dose applied, 12 KJ m^-2^, corresponded to a disease incidence statistically equal to the treatment with 9 KJ m^-2^, but showed a mean lesion diameter twice large (0.008 m) compared to that in treatment with 9 KJ m^-2^ (Fig 1). These results showed a possible damaging effect on melon, where excess of PL can inhibit the fruit defenses against fungal disease, and that the dose of 12 KJ m^-2^ stipulated by the [14] was limiting to the conditions assessed.

Based on these results, we can hypothesize that the PL-treatment with 9 KJ m^-2^ applied here acted as a fungistatic factor inhibiting the mycelial growth of *F. pallidoroseum* and that this behavior is linked to a probable hormetic effect associated with the induction of metabolites synthesis, which possibly activated defense responses against this fungal disease in melon cv. Spanish. For this reason, 9 KJ m^-2^ was chosen for identification of the metabolites produced using a UPLC system and chemometric tools to associate the identification of specific metabolites with disease control mechanism in melon infected with *F. pallidoroseum.*

### Putative metabolite identification by UPLC-QTOF-MS

Initially, infected and uninfected melon samples were evaluated in the positive and negative ionization modes. Finally, the negative ionization mode was chosen because it presents a metabolomic profile with a higher number of compounds at higher intensities, and is more selective and sensitive.

The chemical profile of melon samples was established by analyzing the negative mode (ESI^-^) chromatograms, Fig 2, together with mass spectra. The peaks were numbered according to their elution order, the compounds were tentatively identified by interpretation of their MS and MS/MS spectra determined by QTOF-MS, along with data from literature and open-access mass-spectra databases as MassLynx®. Table 1 lists the MS data of tentatively identified compounds, including experimental and calculated *m/z* for the molecular formula, error, and the fragments obtained by MS/MS, as well as the proposed compound for each peak. In general, 12 metabolites of distinct chemical classes, organic acids, non-protein amino acids, and phenolics, have been tentatively identified.

**Table 1.**
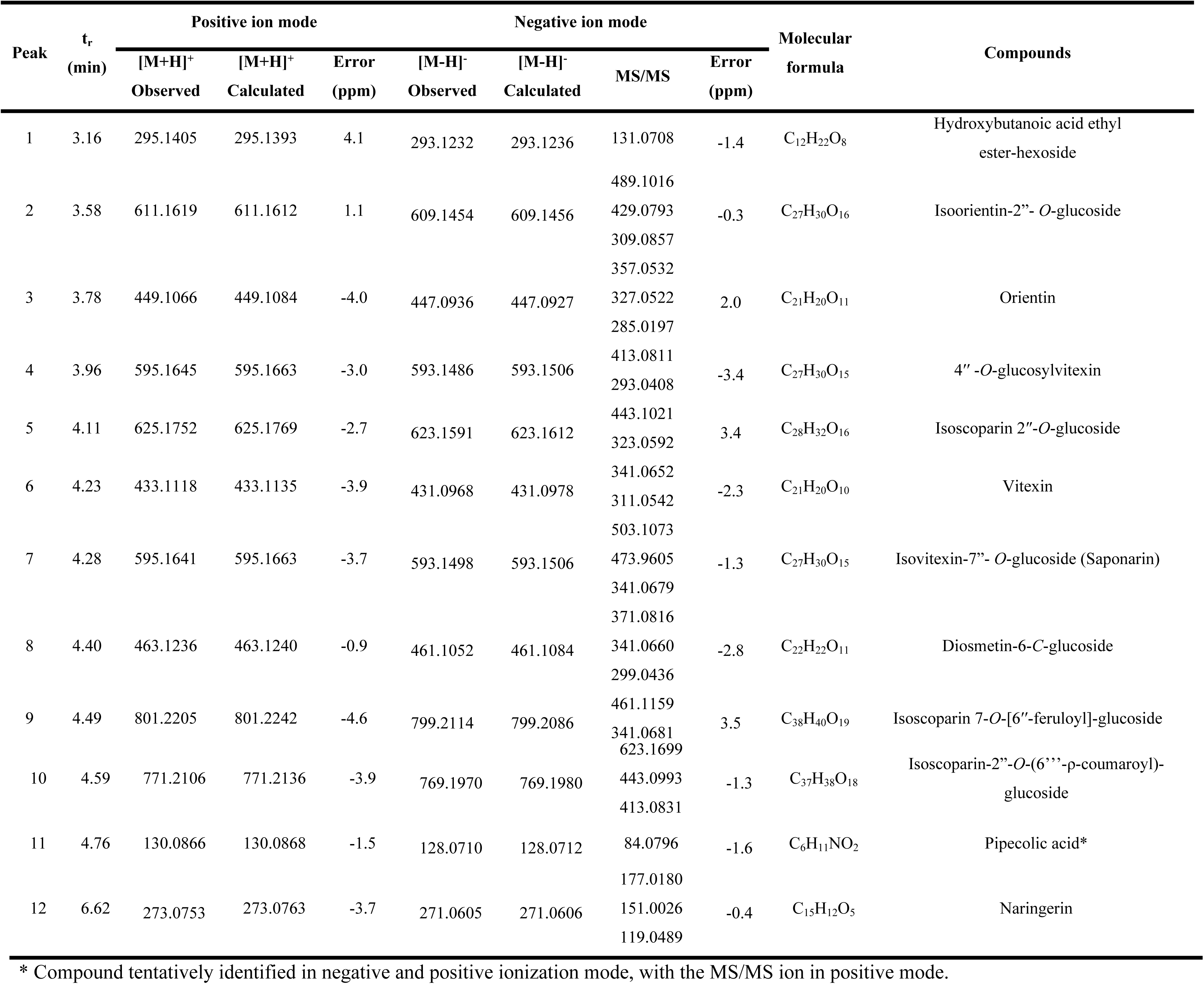
Secondary metabolites tentatively identified by UPLC-QTOF-ESI-MS^E^ in melons treated with PL.

**Fig 2.**
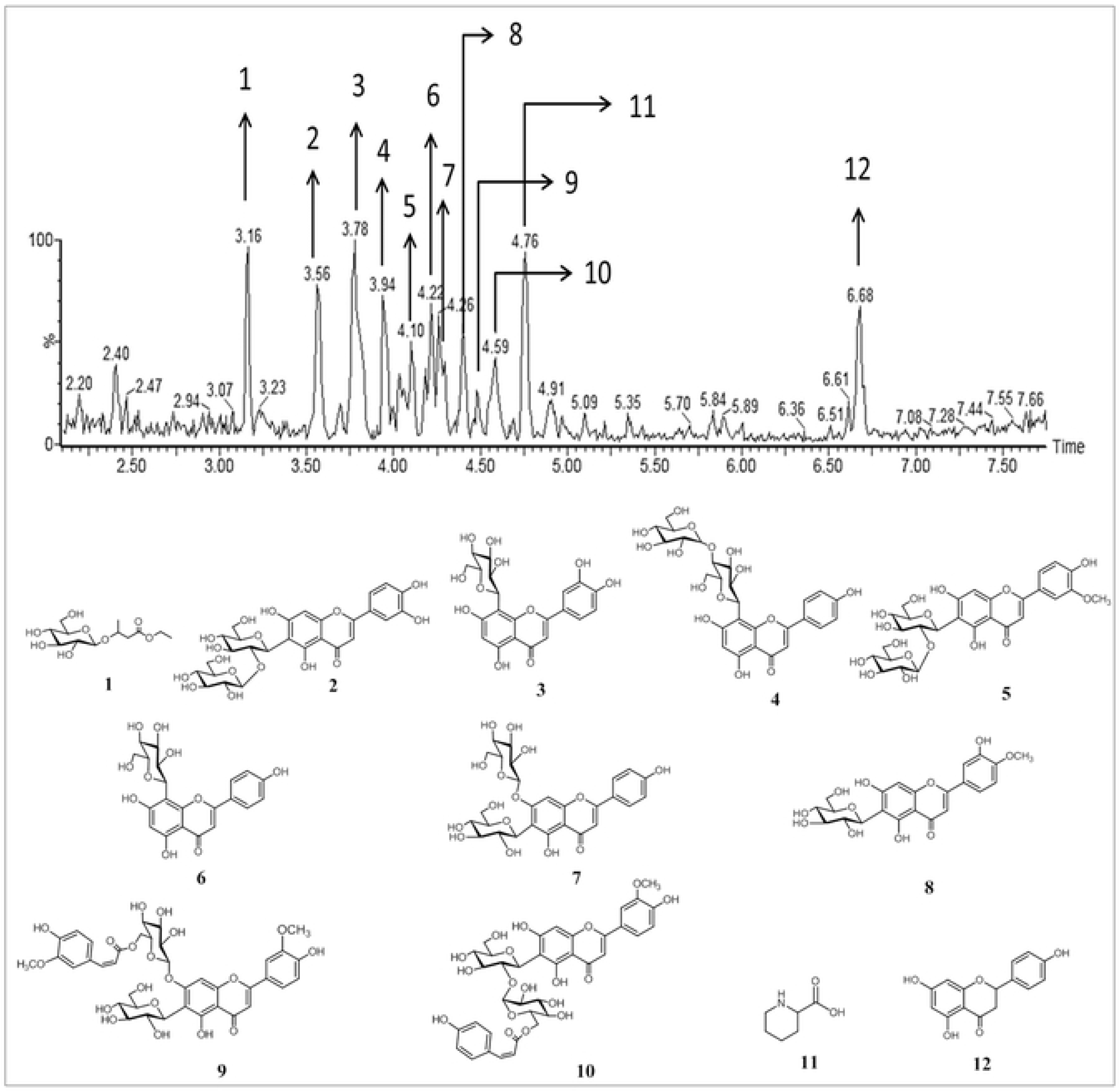
Base-peak chromatogram of melon inoculated with *F. pallidoroseum* and subjected to PL-treatment with 9 KJ m^-2^.

The peak **1** (t_r_ = 3.16 min) was tentatively identified as hydroxybutanoic acid ethyl ester-hexoside. This compound showed the ion *m/z* 293.1232 [M-H]^-^ in MS and the fragment ion *m/z* 131.0708 [M-H-162]^-^ in MS/MS, indicating a pattern loss of the hexoside moiety [27].

The mass spectrum of peak **2** (t_r_ = 3.58 min) showed the precursor ion *m/z* 609.1454 [M-H]^-^. The MS/MS spectrum showed fragment ions *m/z* 489.1016 [M-H-120]^-^, 429.0793 [M-H-180]^-^, and 309.0857 [M-H-180-120]^-^. The losses of 180 and 120 u are significant for diglucosides like sophoroside (1-2 linkages of two glucose molecules). Through the correlation of ions observed in MS and MS/MS, the compound was tentatively identified as luteolin-6-*C*-glucosyl-2”-*O*-glucoside, also known as isoorientin-2”-*O*-glucoside [28].

The mass spectrum of peak **3** (t_r_ = 3.78 min) presents the precursor ion *m/z* 447.0936 [M-H]^-^ that exhibited fragments ions *m/z* 357.0532 [M-H-90]^-^, 327.0522 [M-H-120]^-^, and 285.0197 [M-H-162]^-^. The fragment ion *m/z* 285.0197 [M-H-162]^-^ is the aglycone formed from the loss of the glycosidic group (Table 1 and Fig. 2.) [29]. Thus, the compound was tentatively identified as 8-*C*-glucosyl luteolin, also known as orientin [28].

The compound **4** (t_r_ = 3.96 min) showed in the mass spectrum as the precursor ion *m/z* 593.1486 [M-H]^-^, also exhibiting the fragment ions *m/z* 413.0811 [M-H-180]^-^ and 293.0408 [(aglycone+41)-18)]^-^, which are characteristic of flavone *O*-glucosyl-*C*-glucoside, indicating the presence of sophoroside, and apigenin as aglycone. This compound was characterized as apigenin-6-*C*-glucosyl-2”-*O*-glucoside, also known as 4’’-*O*-glucosylvitexin or isovitexin-2”-*O*-glucoside [28, 30].

The peak **5** (t_r_ = 4.11 min) showed in the mass spectrum a precursor ion *m/z* 623.1591 [M-H]^-^ and its respective fragment ion *m/z* 443.1021 [M-H-162+18]^-^, which suggested loss of a glucose moiety and one unit of water and the fragment ion *m/z* 323.0592 [M-H-120]^-^. These patterns of fragmentation indicate the a presence of a diglucoside linkage and thus, compound **5** was tentatively identified as isoscoparin 2”-*O*-glucoside [31].

The peak **6** (t_r_ = 4.23 min), presented in the mass spectrum the precursor ion *m/z* 431.0968 [M-H]^-^. In the MS/MS spectrum, it showed the fragments ions *m/z* 341.0652 [M-H-90]^-^ and 311.0542 [M-H-120]^-^, indicating the presence of hexose as the monosaccharide and apigenin as aglycone. Therefore, the compound was tentatively identified as 8-*C*-glucosyl apigenin, also known as vitexin [28].

The mass spectrum of peak **7** (t_r_ = 4.28 min) showed the ion *m/z* 593.1498 [M-H]^-^ and the fragment ions *m/z* 503.1073 [M-H-90]^-^ and 473.9605 [M-H-120]^-^. This pattern of loss indicates the presence of a diglucoside similar to that in compound **5** and *m/z* 341.0679 [M-H-120-132]^-^, the loss of 132 Da represents a pentoside. Thus, the compound was tentatively identified as isovitexin-7”-*O*-glucoside, also known as saponarin. [32].

Peak **8** (t_r_ = 4.40 min) in the mass spectrum showed the precursor ion *m/z* 461.1052 [M-H]^-^ and its fragments *m/z* 371.0816 [M-H-90]^-^, 341.0660 [M-H-120]^-^, and *m/z* 299.0436 [M-H-162]^-^. The fragment ion 299.0436 [M-H-162]^-^ represents the aglycone, formed by the loss of a glucoside moiety. Thus, the compound was characterized as diosmetin-6-*C*-glucoside [33]

The peak **9** (t_r_ = 4.49 min) represents compound **9** with a mass spectrum showing the ion *m/z* 799.2114 [M-H]^-^ and its fragment *m/z* 461.1159 [M-H-338]^-^ representing a loss of feruloyl plus a glucoside moiety and *m/z* 341.0681 [M-H-feruloyl-Glucoside-120]^-^.Thus, based on fragmentation, the metabolite was tentatively identified as isoscoparin 7-*O*-[6′′-feruloyl]-glucoside [31]

The peak **10** (t_r_ = 4.59 min) presents in the mass spectrum, the ion *m/z* 769.1970 [M-H]^-^ and its fragments *m/z* 623.1699 [M-H-146]^-^ showing a loss of the coumaroyl moiety; the fragment ion *m/z* 443.0993 [M-H-coumaroyl-162-18]^-^ is the result of successive losses of coumaroyl, glucoside, and a unit of water. The correlation of the ions observed indicates that the metabolite in question is isoscoparin-2”-*O*-(6′′′-*p*-coumaroyl)-glucoside [4]

Negative and positive ionization modes were used to identify pipecolic acid, a non-protein amino acid (homolog of proline). Therefore, the peak **11 (**t_r_ = 4.76 min) showed the ions *m/z* 128.0710 [M-H]^-^ and *m/z* 130.0866 [M+H]^+^ in the MS in the negative and positive ionization modes, respectively. Corroborating with chemical identification, the fragment ion *m/z* 84.0796 [M+H-COOH]^+^ was observed in the positive mode, referring to an aromatic core obtained from the cleavage of the carboxyl group (Table 1 and Fig 2) [34].

In peak **12** (t_r_ = 6.62 min) the ion *m/z* 271.0605 [M-H]^-^ was observed with a fragmentation pattern in MS/MS showing the loss of ring B at *m/z* 177.0180 and by Retro Diels-Alder at *m/z* 151.0026 and *m/z* 119.0489. Thus, it was tentatively identified as flavanone naringenin [35, 36].

### Chemometric analysis

Principal component analysis (PCA) is a multivariate data analysis method that can synthesize data from an original matrix with many variables in the set of smaller orthogonal variables [37, 38]. We confirm that the chemometric analyses were centered in PL-treatment with a dose of 9 KJ m^-2^, considered here as treatment for disease control in melon inoculated with *F. pallidoserum* (Fig 1).

Therefore, PCA-2D was performed to discriminate different treatments groups according to their metabolic profiles represented by retention time and the mass-to-charge ratio (rt-*m/z*) from the UPLC-QTOF-MS^E^ analysis (Fig 3). The PCA-2D showed perfect separation of all groups evaluated with 89% of the total cumulative variance in the diaxial axes PC1 and PC2 (R^2^X[1] = 0.7401 and R^2^X[2] = 0.1571), with a data noise level of 6%, showing a robust model regarding data certainty. The formation of groups was related to the similarity between biological triplicates.

**Fig. 3.**
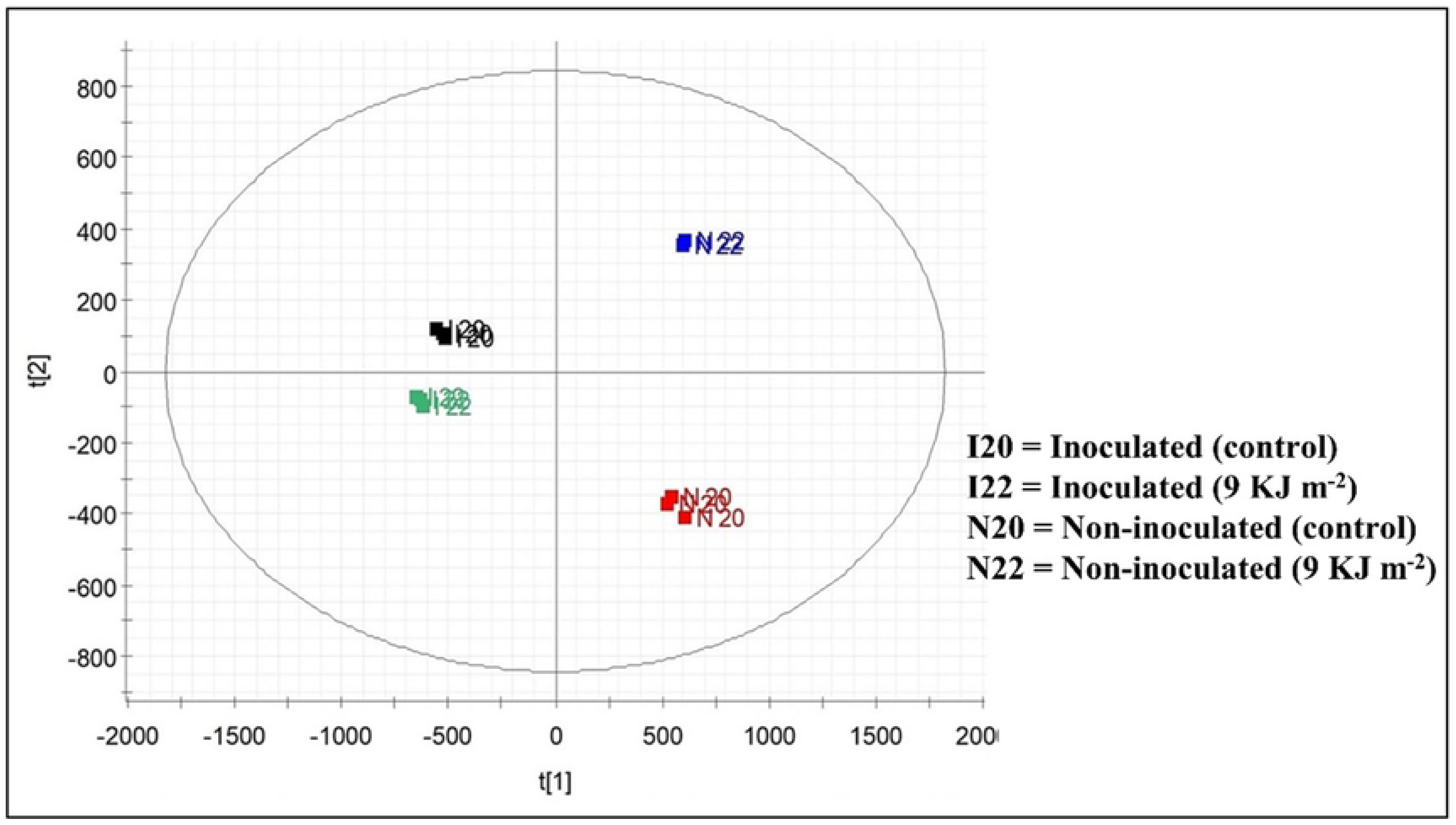
Discrimination of treatment groups by PCA-2D.

The PCA-2D in the PC1 showed the inoculated group (negative scores) and non-inoculated group (positive scores); whereas in the PC2 (positive scores), discrimination of the inoculated control and non-PL-treated group was observed. PC2 (negative scores) also showed the separation of inoculated PL-treated and non-inoculated groups (Fig 3). The separation of negative and positive groups in PC1 and PC2 is clearly linked to the differences of each metabolomic profile [39]. For this reason, OPLS-DA chemometric analysis was applied to the UPLC-QTOF-MS^E^ data to compare samples according to metabolites that influenced on disease control against *F. pallidoroseum* in melons treated with PL. OPLS-DA is an analysis method used to study ions that contribute to experimental sample classification, and the classification between two groups in the OPLS-DA model can be visualized in the form of a score chart and scatter plot (S-plot).

Fig 4A summarizes the separation between non-inoculated and non-treated fruit (control) and non-inoculated fruit treated with PL (9 KJ m^-2^), with respect to metabolic responses in the function of PL as abiotic stress, through OPLS-DA (R^2^X(cum) = 0.9232). The S-Plot generated from OPLS-DA, with VIP > 1.0 and *p* < 0.05 showed potential biomarkers between the treatments evaluated.

**Fig 4.**
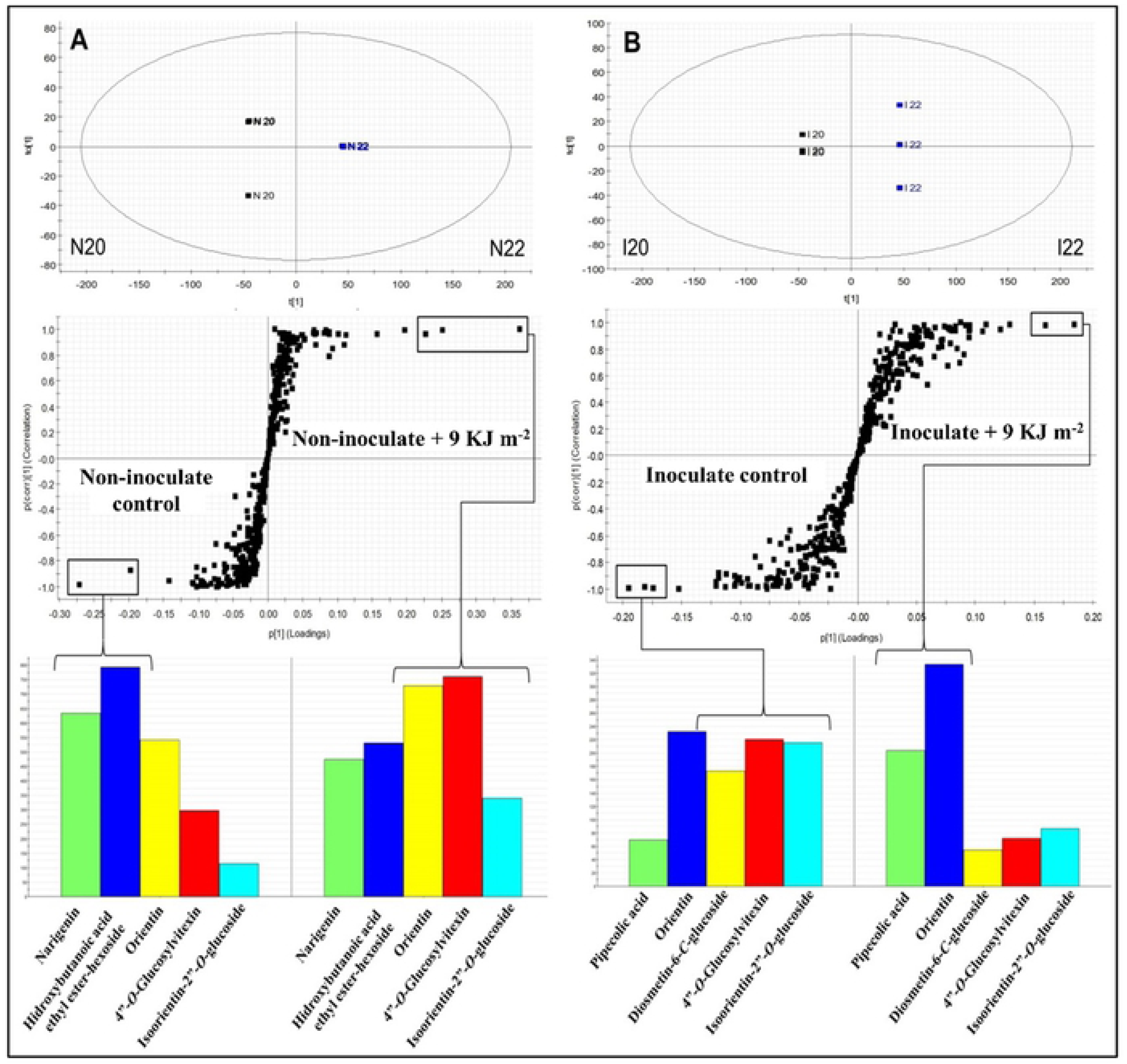
OPLS-DA, S-Plot graphs, and intensity of biomarkers between non-inoculated control and non-inoculated and PL-treated melons (9 KJ m^-2^) (**A**). OPLS-DA, S-Plot graphs, and intensity of markers between inoculated control and inoculated and PL-treated melons (9 KJ m^-2^) (**B**).

The control group showed synthesis of principal secondary metabolites such as naringenin (peak **12**), hydroxybutanoic acid ethyl ester-hexoside (**1**), isoorientin-2”-*O*-glucoside (**2**), orientin (**3**), and 4”-*O*-glucosylvitexin (**4**). However, an upregulation of naringenin (**12**) and hydroxybutanoic acid ethyl ester-hexoside (**1**) was highlighted (Fig 4A). Naringenin is a flavonoid that serves as a primer for more advanced flavonoid structures and as a substrate for glycosylation reactions. Hydroxybutanoic acid ethyl ester-hexoside is a compound has been previously described in other varieties of melon such as Piel de Sapo, Galia, and Cantaloupe [27]. Interestingly, naringenin (**12**) and hydroxybutanoic acid ethyl ester-hexoside (**1**) were downregulated in melons treated with hormetic PL-radiation, whereas specific flavonoid compounds such as orientin (**3**), 4”-*O*-glucosylvitexin (**4**), and isoorientin-2”-*O*-glucoside (**2**) were up-regulated (Fig 4A), thus these compound which were up-regulated might be considered as biomarkers caused by PL-treatment. Accumulation of flavonoid precursors such as phenylalanine ammonia lyase into vacuoles present in the epidermal and subepidermal mesophyll tissues of fruit stimulate plant defense mechanisms under specific PL-radiation conditions [24]. Many phenylpropanoids have been associated with induced disease resistance and disease control. According to Jung et al. (2013) the flavonoids orientin, isoorientin-2”-*O*-glucoside, and 4”-*O*-glucosylvitexin are secondary metabolites involved in antioxidant activities against abiotic stress in rice leaves (*Oryza sativa* cv. Ilmi) exposed to different conditions of LED light radiation.

The differences in inoculated (control) and inoculated plus PL-treated (9 KJ m^-2^) melons are shown in Fig 4B. Separation of the two treatments groups by the OPLS-DA graph (R^2^X(cum) = 0.8882) indicated the differences among groups according to their chemical profiles. The S-Plot obtained from the OPLS-DA graph, demonstrated different potential biomarkers between groups for VIP > 1.0. Inoculation of *F. pallidoroseum* mainly induced the synthesis of glycosylated flavonoids such as diosmetin-6-*C*-glucoside (**8**), isoorientin-2”-*O*-glucoside (**2**), and 4’’-*O*-glucosylvitexin (**4**) (Fig 4B), and this behavior is due to the presence of flavonoids at the infection site that are responsible for defense mechanisms against pathogens [40]. However, these metabolites which are a natural response of fruit against de pathogen were not sufficient to control the growth pathogen as shown by control treatment at Figure 1.

Biotic and abiotic stress might regulate systemically the defense mechanism by induce systemic resistance (ISR) or systemic acquired resistance (SAR) [41, 42]. PL-treatment and inoculation with *F. pallidoroseum* led by SAR to the upregulation of two major biomarkers in melon, pipecolic acid (**11**) and orientin (**3**) (Fig 4B), indicating a change in the metabolic pathway, where the fruit preferentially used these specific compounds as phytochemical markers in response to the treatment [43]. The presence of pipecolic acid was verified in both groups that were inoculated with *F. pallidoroseum*. The presence of pipecolic acid in pathogen inoculation sites is associated with plant defense responses and acts as a regulator of inducible plant immunity [44]. Thus, when the PL-treatment is applied the pipecolic acid together with orientin is up-regulated significantly achieving disease control (fungistatic effect) which also seen at Figure 1 on treatment 9 KJ.m^2^. Curiously, pipecolic acid is the supposed precursor of betaines that are accumulated in the cytoplasm as a result of physiological plant responses to stress phenomena induced in a large part by adverse environmental conditions. These compounds that are biochemically inert in the cell are synthesized from some specific amino acids such as serine, alanine, methionine, the non-protein amino acid γ-aminobutyric acid, and some cyclic amino acids such as proline and pipecolic acid. Biosynthesis of betaines in the cytoplasm under abiotic stress conditions is mainly due to the action of methyltransferases, which utilize S-adenosylmethionine as a methyl group donor [43].

Fig 5A shows the results of the OPLS-DA graph (R^2^X(cum) = 0.9684) between the inoculated and non-inoculated groups in the function of *F. pallidoroseum* as a biotic stress. The S-Plot originating from OPLS-DA data demonstrated potential biomarkers in each group, with values for VIP > 1.0 and *p* < 0.05.

**Fig 5.**
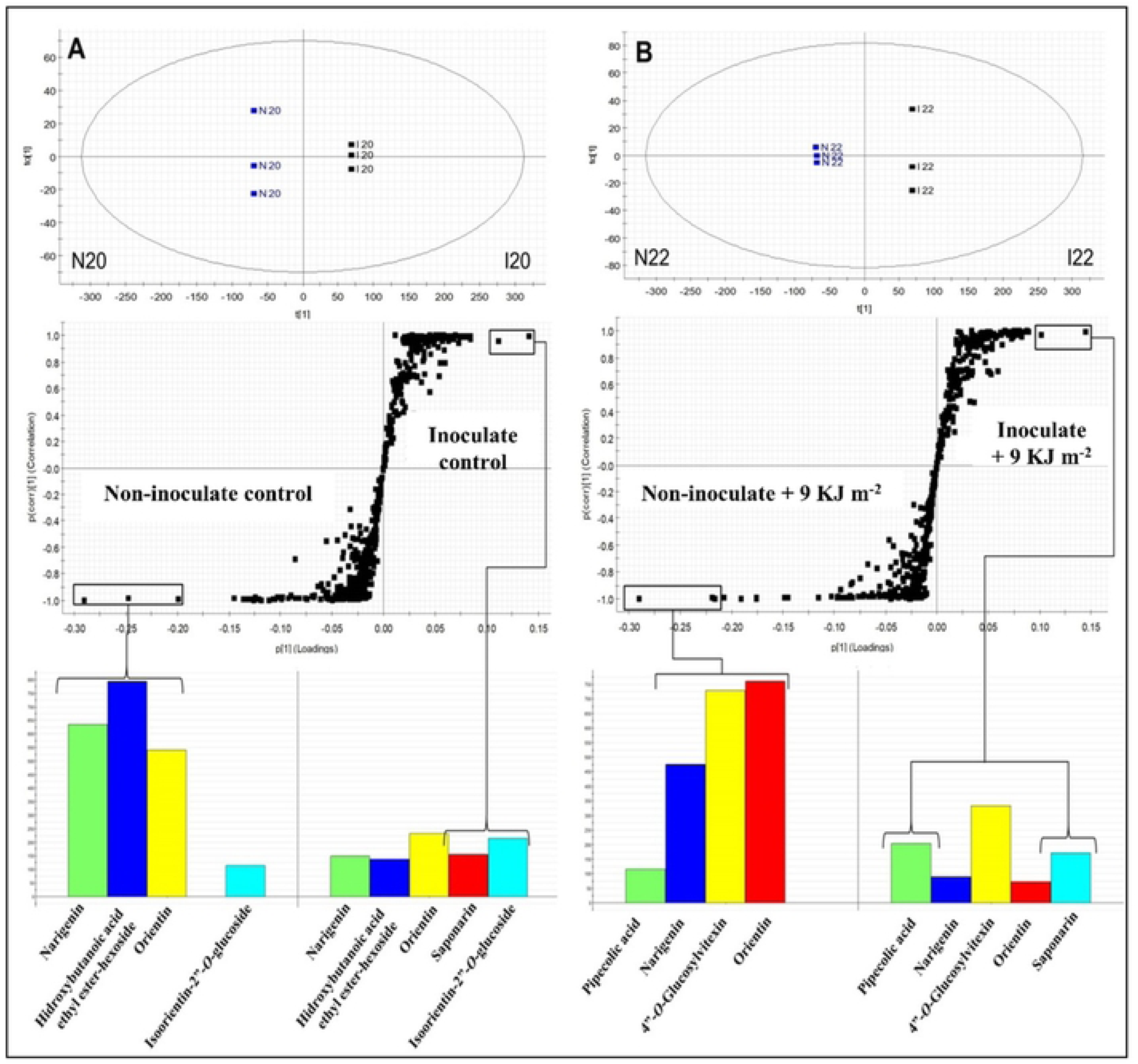
OPLS-DA, S-Plot graphs, and intensity of biomarkers between non-inoculated and inoculated melons (**A**). OPLS-DA, S-Plot graphs, and intensity of markers between treated non-inoculated (9 KJ m^-2^) and PL-treated inoculated melons (9 KJ m^-2^) (**B**).

The number of compounds up-regulated in non-inoculated melons is much higher when that compared to inoculated melons, highlighting naringenin (**12**), hydroxybutanoic acid ethyl ester-hexoside (**1**) and orientin (**3**) (Fig 5A). However, saponarin (**7**) was a biomarker present in group inoculated control melons and isoorientin-2”-*O*-glucoside (**2**) shows a little up-regulation on the same group (Fig 5A). Hydroxybutanoic acid ethyl ester-hexoside is a compound linked to amino acid groups and findings have indicated significant interconnections between different branches of amino acid metabolism and plant resistance to pathogens [45, 46]. Naringenin is a flavonoid showing metabolic activity associated with barley resistance to *Fusarium graminearum* [47].

The results observed in Fig. 5B show the separation of distinct groups, where the OPLS-DA plot (R^2^X(cum) = 0.9798) presents differences between the non-inoculated fruit and inoculated plus PL-treatment. The S-Plot obtained by the OPLS-DA graph points out the potential markers for each group. Upregulation of flavonoids such as naringenin (**12**), orientin (**3**) and 4”-*O*-glucosylvitexin (**4**) in non-inoculated PL-treated fruit (9 KJ m^-2^) is much higher when compared to the inoculated and PL-treated (9 KJ m^-2^) group (Fig. 5B). The synthesis of flavonoids in detriment to abiotic stress observed in PL treatment occurs due to a change in the metabolic pathway of fruit in response to the medium [48]. On the other hand, inoculated and PL-treated melons accumulated a glycosylated flavonoid identified as saponarin (**7**), and pipecolic acid (**11**) as biomarkers [49]. Nevertheless, the presence of pipecolic acid on non-inoculated and PL-treated (9 KJ m^-2^) group (Fig 5B) shows that this metabolite might be synthetized by abiotic stress also. Saponarin is an antioxidant belonging to the flavones and is to inhibiting malonaldehyde formation in barley. In a normal reaction, malonaldehyde is formed from oxidized lipids on the skin surface in barley leaves by UV irradiation [50]. The effect of PL treatment in melons inoculated with fungi resulted in a response mediated by the synthesis of pipecolic acid in the cellular medium, which resulted in increased levels of betaines in the cell, that induce the fruit immunity system.

## Conclusions

In the present study, a PL-dose of 9 KJ m^-2^ in melon inoculated with *Fusarium pallidoroseum* controlled the disease promoted by this pathogen (fungistatic effect), and induced metabolic variation in the fruit defense system. Pipecolic acid (**11**) and orientin (**3**) were the two major biomarkers associated with postharvest disease control against *F. pallidoroseum* in infected melons treated with a pulsed radiation. This study also showed that fruit subjected to biotic and abiotic stresses, separately, demonstrated different metabolic responses, with a chemical profile in response to each stress. In addiction the compounds orientin (**3**), 4”-*O*-glucosylvitexin (**4**), and isoorientin-2”-*O*-glucoside (**2**) are the metabolites in response only PL-treatment. Our findings highlight that the application of PL technology provided control against the postharvest disease of melon cv. Spanish by directly controlling the growth of *F. pallidoroseum* through the synthesis/up-regulation of specific compounds that acted as principal biomarkers of the defense system against pathogen. Thus, PL can be readily proposed as a new postharvest technology alternative to chemical fungicides and may be applied to reduce the delay caused by *F. pallidoroseum* in melons var. Spanish during their export and storage.

## Acknowledgments

The authors are gratefully acknowledged for support from the Brazilian Company of Agricultural Research (EMBRAPA), National Council for Scientific and Technological Development (CNPq), National Institute of Science and Technology (INCT BioNat, Brazil) and commercial growing field of Norfruit Northeast Fruit.

